# TLR3 deficiency exacerbates the loss of epithelial barrier function during genital tract *Chlamydia muridarum* infection

**DOI:** 10.1101/459636

**Authors:** Ramesh Kumar, Haoli Gong, Luyao Liu, Nicole Ramos-Solis, Cheikh I. Seye, Wilbert A. Derbigny

## Abstract

**Problem:** *Chlamydia trachomatis* infections are often associated with acute syndromes including cervicitis, urethritis, and endometritis, which can lead to chronic sequalae such as pelvic inflammatory disease (PID), chronic pelvic pain, ectopic pregnancy, and tubal infertility. As epithelial cells are the major cell type productively infected during genital tract *Chlamydia* infections, we investigated whether *Chlamydia* has any impact on the integrity of the host epithelial barrier as a possible mechanism to facilitate the dissemination of infection, and examined whether TLR3 function modulates its impact.

**Method of Study:** We used wild-type and TLR3-deficient murine oviduct epithelial (OE) cells to ascertain whether *C. muridarum* infection had any effect on the epithelial barrier integrity of these cells as measured by transepithelial resistance (TER) and cell permeability assays. We next assessed whether infection impacted the transcription and protein function of the cellular tight-junction (TJ) genes for claudins1-4, ZO-1, JAM1 and occludin via quantitative real-time PCR (qPCR) and western blot.

**Results:** qPCR, immunoblotting, transwell permeability assays, and TER studies show that *Chlamydia* compromises cellular TJ function throughout the course of infection in murine OE cells, and that TLR3 deficiency significantly exacerbates this effect.

**Conclusion:** Our data show that TLR3 plays a role in modulating epithelial barrier function during *Chlamydia* infection of epithelial cells lining the genital tract. This proposes a role for TLR3 signaling in maintaining the integrity of epithelial barrier function during genital tract *Chlamydia* infection, a function that we hypothesize is important in helping limit the chlamydial spread and subsequent genital tract pathology.

## Introduction

*Chlamydia trachomatis* is a gram-negative intracellular bacterium and the cause of the disease chlamydia, which is the most common sexually transmitted infection in the United States, with over 1.7 million cases reported in the US in 2017 alone[1]. Genital tract infections with *C. trachomatis* are associated with many acute syndromes including cervicitis, urethritis, and endometritis [2]. Complications from chronic infections include pelvic inflammatory disease (PID) and its sequelae of chronic pelvic pain, ectopic pregnancy, and tubal infertility [3]. Although *Chlamydia* is treatable with antibiotics, infected individuals are often asymptomatic; which facilitates the spread of the bacterium through further sexual contact. As a result, *Chlamydia* infections have continued to rise despite the implementation of screening and early intervention strategies [4]. The ultimate goal in developing more effective therapeutic measures against *Chlamydia* infection is to identify aspects of host immunity that will augment clearance, while minimizing immune responses that lead to genital tract pathology.

As an obligate intracellular pathogen, Chlamydiae are known to interact with host-cell pattern recognition receptors (PRRs), including a variety of intracellular cytosolic receptors and Toll-like receptors (TLRs) [5–10]. TLRs are PRRs that recognize conserved microbial molecules or pathogen-associated molecular patterns (PAMPs) [11]. Stimulation of TLRs by chlamydial PAMPs triggers cytokine responses critical to the establishment of innate and adaptive immune responses [5, 7, 12–15]. It is critically important to identify the TLRs that induce the specific inflammatory mediators that cause scarring and fibrosis, and define therapeutic approaches to prevent this process.

TLR3 is a receptor for double-stranded RNA (dsRNA) and is known to activate transcription of IFN-β via the adaptor protein Toll-IL-1 receptor (TIR) domain-containing adaptor molecule-1 (TICAM-1) [also called TIR-domain-containing adapter inducing IFN-β (TRIF)] [16, 17]. TLR3 is expressed intracellularly and on the cell surface on human fibroblasts [17]; however, TLR3 has exclusive intracellular expression in most other cell types [18–20]. TLR3 has been identified as the major MyD88-independent PRR stimulated in the type-1 IFN responses to many different viral infections due to its intracellular localization [21–26]. Conversely, its role in bacterial infection is poorly understood, particularly since bacteria are not known to possess a dsRNA moiety. We previously showed that *C. muridarum* infected murine oviduct epithelial (OE) cells secrete IFN-β in a mostly TLR3 dependent manner, and that they demonstrate dramatic reductions in the syntheses of other inflammatory immune mediators in addition to IFN-β [6, 8]. Results from our recent *in vivo* study show that TLR3-deficient mice have significantly different levels of several key innate-immune factors secreted into their genital tracts during *C. muridarum* infection, and demonstrate an altered recruitment of CD4^+^ T-cells to the reproductive tract when compared to wild-type control mice [27]. Because of this altered immune response to *C. muridarum* infection, we routinely observe higher bacterial burdens early and mid-infection, and more substantial reproductive tract pathology in the TLR3-deficient mice. These findings have proved our hypothesis that a less than optimal immune response in the TLR3-deficient mice increases the severity of *Chlamydia* infection, and provides an impetus to investigate putative mechanisms that are invoked to elicit the more progressive disease in the TLR3-deficient mice.

In this study, we further delineate the role of TLR3 in genital tract pathology associated with *Chlamydia* infection, by examining the impact that TLR3 signaling has on bolstering the host’s protective epithelial barrier that is designed to limit the spread of infection. Here, we show that show that *Chlamydia* has a definite impact on cellular TJ function throughout the course of infection in murine OE cells, and that TLR3 function significantly modulates this effect.

## Materials and Methods

### Ethics Statement

C57BL/6J (control) and TLR3-deficient mice were purchased from The Jackson Laboratory (Bar Harbor, ME) at 6-8 weeks of age. All mice were provided food and water *ad libitum*, and kept on a standard 12-hour light/dark cycle. All mice were given at least a 1-week period where they were allowed to acclimate to their new environment. After acclimation, female C57BL/6NJ and TLR3-deficient mice were injected with 2.5 mg Depo-Provera after being briefly anesthetized with isoflurane prior to any experiment, and all mice were allowed to recover for no less than 1-week after the Depo-Provera treatment. To alleviate any possible distress, the mice were also briefly anesthetized with isoflurane prior to either intravaginal infection with *C. muridarum*, the insertion of calcium alginate swabs, or the insertion of aseptic vaginal sponges. All mice were monitored daily for lethargy, signs of vaginal bleeding, and/ or death. None of the mice exhibited morbidity during the course of these studies. All mice were euthanized by either exposure to isoflurane or inhalation of carbon dioxide, followed by exsanguination. The Indiana University Institutional Animal Care and Use Committee (IACUC) approved all experimental animal protocols. All care was used to ensure that steps were taken to ameliorate animal suffering in all work involved in the removal of genital-tract tissue.

### Cells and bacteria

The cloned oviduct epithelial cell lines OE-TLR3(-) and OE-129WT [6] were grown at 37°C in a 5% CO2 humidified incubator and maintained in epithelial-cell media: 1:1 Dulbecco’s modified Eagle medium:F12K (Sigma-Aldrich, St. Louis, MO), supplemented with 10% characterized fetal bovine serum (FBS) (HyClone), 2 mM L-alanyl-L-glutamine (Glutamax I; Gibco/ Invitrogen, Carlsbad, CA), 5μg of bovine insulin/ ml, and 12.5ηg/ ml of recombinant human FGF-7 (keratinocyte growth factor; Sigma) as previously described [6, 28].

*Mycoplasma*-free *C. muridarum* Nigg, previously known as *C. trachomatis* strain (MoPn), was grown and titrated in McCoy cells (ATCC) as described [28, 29]. Elementary bodies (EBs) were harvested from infected cells, resuspended in SPG buffer (250mM sucrose, 10mM sodium phosphate, and 5mM L-glutamic acid, pH 7.2), and quantified on McCoy cells using methodology described previously [8, 28, 30].

### Mice and infections

Wild-type control mice C57BL/6NJ [Stock No 005304] and TLR3-deficient mice B6N.129S1-Tlr3tm1Flv/J [Stock No 009675] were purchased from The Jackson Laboratory (Bar Harbor, ME) at 6-8 weeks of age. All mice were housed in Indiana University Purdue University-Indianapolis specific pathogen-free (SPF) facilities. Age-matched mice were used at 14–16 weeks for the experiments in this study. The Indiana University Institutional Animal Care and Utilization Committee (IACUC) approved all experimental protocols.

Infections of mice were done as described in [5] with some minor modifications. Groups of 6 mice (1 mock treated; 5 infected) were treated with 2.5mg of Depo-Provera (medroxyprogesterone acetate; Pfizer; New York, NY) in 0.1 ml saline one week before vaginal infection with 10^5^ IFU *C. muridarum* (approximately 100 times the ID50) in 10μl sucrose-phosphate-glutamic acid (SPG) buffer (250mM sucrose, 10mM sodium phosphate, and 5mM L-glutamic acid, pH 7.2). To procure genital tract tissue from the infected mice, we used methodology described in [31] with some modification. Briefly, the entire genital tract was sterilely harvested from each mouse on either day 3, day 5, or day 7 post-infection. The segment encompassing the oviduct and uterine horn was carefully excised from each side of the genital tract, and both sides from the same mouse were combined to comprise the entire upper genital tract (UGT) tissue sample from that particular mouse. Tissue samples were quickly frozen at −80°C until time of use.

*In vitro* infection of the oviduct epithelial cell lines OE129WT and OE129TLR3(-/-)C19 were performed as described in [6, 32]. Briefly, the OE cells were plated in their respective cell-culture plates and were infected when confluent. The cells were infected with an inoculum containing anywhere from 1 to 10 inclusion forming-units (IFU) of *C. muridarum*/ cell (depending upon the experiment), and the cell-culture plates were centrifuged 1000 x *g* for 1hr at room temperature in a table-top centrifuge to synchronize infection prior to being returned to the 37°C CO2 incubator.

### Measuring *C. muridarum* organism recovery from upper genital tract tissue

We used a similar method to monitor the upper genital tract infection in mice as described elsewhere [31]. To recover viable *Chlamydia* from the UGT of *C. muridarum* infected wild-type and TLR3-deficient mice, each mouse’s UGT tissue sample was homogenized in 300μl of SPG using a 2-ml Dounce tissue grinder (Sigma) and subsequently passaged through a 20-gauge syringe. After a brief sonication, the released chlamydial EBs were titrated on McCoy cell monolayers as described previously in [8, 28, 30]. Briefly, chlamydial elementary bodies harvested from infected tissue were serial diluted 1:10 in SPG, and quantified as inclusion forming units (IFUs) per milliliter on McCoy cell monolayers. The total number of IFUs per ml was calculated based on enumeration of fluorescent inclusion bodies in the cells whereby antibody specific for chlamydial LPS was used to detect chlamydial inclusions in the infected McCoy cells. Detection of the chlamydial LPS was done via Alexa Fluor 488 anti-mouse IgG secondary antibody (Invitrogen/Life Technologies; Carlsbad, CA), and immuno-staining results were scanned and recorded by EVOS imaging system (Thermo-Fisher, Pittsburgh, PA).

### Electric Cell-Substrate Impedance Sensing (ECIS)

Trans-epithelial resistance (TER) of *C. muridarum* infected OE cell monolayers were measured with the ECIS system using the Zθ (Theta) model and using the 8W10E+ (8-well) arrays (Applied Biophysics Inc; Troy, NY). Prior to seeding the cells, the array chambers were treated with a 10mM solution of cysteine for 10 min at room temperature to stabilize the electrode. After removing the stabilization solution, the wells were washed twice with sterile distilled water, and seeded with either the OE-129WT cells or the OE-TLR3(-) cells at a density of 10^5^ cells/cm^2^ in 400ml total volume epithelial media to achieve rapid confluence. After a brief centrifugation for 10 min at 1000 x *g* to attach cells to the surface, the cells were incubated at 37°C in a CO_2_ incubator for 4hrs to become completely confluent, which was verified by microscopy prior to proceeding to the next step. After 4hrs of incubation, IFN-β was added to certain wells containing OE-TLR3(-) cells to achieve a final concentration of 50U/ml and returned to the CO_2_ incubator for an additional 1hr. After the 1hr incubation with IFN-β, all cells were infected with a multiplicity of infection (MOI) of 1 IFU/cell using the method described above. Resistance, capacitance, and impedance information was collected continuously for 48hrs post-infection. Each experiment had two replicate wells per array and the experiment was repeated at least six times.

### Transwell cell permeability assays

OE-129WT and OE-TLR3(-) cells were seeded at a density of 10^5^ cells/cm^2^ on the TC-coated polyester membrane inserts from the Corning™ Transwell™ 24-well plate system (Costar; Brumath, France). IFN-β pre-treatment and infection with an MOI of 1 IFU/cell *C. muridarum* was performed as described above after cells had been growing in the CO^2^ incubator for 6hrs after seeding. FITC-dextran (molecular mass of 70kDa; Sigma) was used as an index of macromolecular permeabilization: Briefly, FITC-dextran was added to epithelial media to a final concentration of 1mg/mL in 400ul that was added to the upper chambers of the Transwell system. The lower chamber contained 300ul epithelial media and 150uL samples were taken from the lower chamber at 6-hour intervals, and the same volume of epithelial medium was replaced in this chamber to prevent fluid movement caused by hydrostatic pressure. The fluorescence was measured with a BioTek FLx800 fluorimeter (BioTek; Winooski, VT) using 480nm and 520nm as the excitation and emission wavelengths, respectively.

### RNA purification and quantitative real-time polymerase chain reaction (qPCR)

OE129-WT cells, OE-TLR3(-) cells, and OE-TLR3(-) cells that pre-treated with 50U/ml recombinant IFN-β were grown to confluence before being either mock-infected or infected with 10 IFU/ cell *C. muridarum*. The OE-TLR3(-) cells pre-treated with IFN-β had the 50U/ ml IFN-β added to the media 1hr prior to infection as described in [8]. Cell lysates were harvested at either 0, 8, 12, 20, or 36hr post-infection, and total cell RNA was isolated using the RNeasy plus kit (Qiagen; Valencia, CA) according to manufacturer’s protocol. The DNA-free RNA samples were quantified using the NanoDrop spectrophotometer (Thermo Scientific), and cDNA was obtained with the Applied Biosystem’s high-capacity cDNA reverse transcription kit (Thermo Fisher) using 500ng total cell RNA. Target gene cDNA was amplified using Applied Biosystem’s TaqMan gene-expression master kit in reactions containing primers for the various tight-junction (TJ) genes and/or the β-actin control primers (Table 1) according to manufacturer’s protocol. Quantitative measurements were performed via ABI7500 real time PCR detection system (Thermo Fisher). Relative expression levels were measured as a fold increase in mRNA expression versus mock controls and calculated using the formula 2^−ΔΔCt^ as described in [33].

**Table 1.**
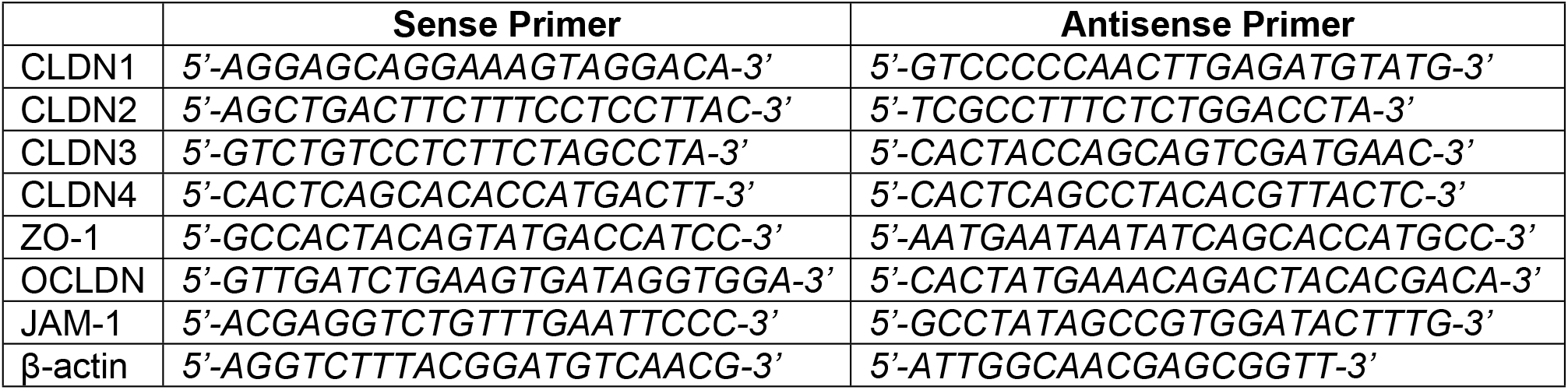
Primers for qPCR

### SDS PAGE and Western blotting

OE-129WT cells, OE-TLR3(-) cells, and OE-TLR3(-) cells that pre-treated with 50U/ml recombinant IFN-β were grown to confluence before being either mock-infected or infected with 10 IFU/ cell *C. muridarum*. Cell lysates were harvested at either 0, 8, 12, 20, or 36hr post-infection, and were analyzed for tight-junction protein expression by SDS-PAGE and subsequent Western blotting techniques as described previously [32]. In brief, equal protein amounts were separated on 4-12% Tris-Glycine Gel (Novex, Invitrogen Carlsbad CA, USA) and blotted onto PVDF membrane (Amersham Hybond™, GE Healthcare Life Sciences; Chicago, Illinois, USA). After blocking, PVDF membranes were immunoblotted with primary antibodies: rabbit anti-mouse claudin-1 (1:200 dilution; catalog# 51-9000), rabbit anti-mouse claudin-2 (1:100 dilution; catalog# 51-6100), rabbit anti-mouse claudin-3 (1:50 dilution; catalog# PA5-16867), mouse anti-mouse claudin-4 (1:50 dilution; catalog# 32-9400), rabbit anti-mouse ZO-1(1:2000 dilution; catalog# 61-7300) and goat anti-mouse JAM1 (1:5000 dilution; catalog# PA5-47059), all purchased from Invitrogen. Rabbit anti-mouse occludin (1:250 dilution; catalog# LS-B2187) was purchased from LsBio (LifeSpan BioSciences Inc., Seattle, WA) and the monoclonal antibody specific for mouse β-actin (1:50000 dilution; catalog# A1978) antibody was purchased from Sigma. The HRP conjugated goat anti-rabbit IgG (catalog# 32460) and goat anti-mouse IgG (catalog# 32430) were purchased from Thermo Scientific, while the chicken anti-goat IgG (catalog# CkxGt-003-EHRPX) was purchased from ImmunoReagent Inc. (Raleigh, NC). All HRP-conjugated antibodies were diluted to 1:7500 dilution and served as secondary antibodies in this study. Detection of specific proteins was done using Super-Signal West-Dura extended duration substrate (Thermo Scientific), and was carried out according to manufacturer’s protocol. Protein bands were quantified by NIH software Image-J, freely available online. The relative band intensity of each tight-junction protein was obtained by the normalization to the band intensity of loading control β-actin. Experiments were repeated at least three times.

### Immunofluorescent staining of tight junction proteins

OE-129WT cells and OE-TLR3(-) cells were grown to confluence in 96-well Ibidi plates (Ibidi USA; Fitchburg, Wisconsin) before being either mock-infected or infected with 1 IFU/ cell *C. muridarum* at 37°C for up to 36hrs. At the specified time-point, the cells were carefully rinsed with PBS before being fixed 10 minutes at room temperature using 10% neutral-buffered formalin (Sigma). The monolayers were washed at room temperature 3 times for 5 minutes using PBS. After washing in PBS, the cells were blocked and permeabilized for 30 min using PBS containing 1% BSA and 0.1% saponin. After permeabilization, cells were washed in blocking buffer (PBS + 1% BSA) at room temperature for 5 min, and the primary antibody (either Claudin-1; 1:50 dilution, ZO-1; 1:100 dilution, or JAM-1; 1:200 dilution) was added to the cells for 1hr at room temperature. After washing at room temperature 3 times for 5 minutes using PBS, the cells were then incubated in the dark at room temperature for 1hr using either goat anti-rabbit, goat anti-mouse, or donkey anti-goat 2° antibody conjugated with Alexa fluor-594 (1:1000 dilution, Molecular Probes). Finally, the cells were washed 3 times for 5 minutes using PBS and counterstained with DAPI to identify the nuclear DNA. Duplicates processed without primary antibodies served as negative controls. Fluorescence was imaged using a Leica DMI 6000B inverted fluorescent microscope.

### Statistical Analysis

Numerical data are presented as mean ± (SD). All experiments were repeated at least three times, and statistical significance was determined using Student’s two-tailed *t*-test. 2-way *ANOVA* analyses in Graphpad Prism were used to analyze the differences in IFUs recovered from mouse upper genital tract tissue homogenates. Statistically significant differences are shown by asterisks (*), and with the minimum criteria being *p*<0.05.

## Results

### *Chlamydia muridarum* ascends more rapidly into the upper genital tract tissues of TLR3-deficient mice

We previously showed that TLR3-deficient mice exhibited significantly higher levels of chlamydial shedding from the lower genital tract when compared to wild-type (WT) mice during the first four weeks of infection [6, 27]. Additionally, our earlier *in vitro* studies showed that *Chlamydia* replication in oviduct epithelial (OE) cells deficient in TLR3 was more robust than in the wild-type OE cells [8]. The data from those previous experiments showed that TLR3 had a biological impact on the innate immune response to *Chlamydia* infection in mice, and we hypothesized that TLR3 deficiency may have subsequent impact on *Chlamydia’s* ability to spread and ascend the female genital tract. To ascertain whether TLR3’s function indeed has a role in the chlamydial spread and ascension into the upper genital tract (UGT) tissue of female mice, we infected WT and TLR3 deficient mice with 10^5^ IFU *C. muridarum*, and harvested the UGT tissue from mice at either day 3, 5, or 7 post-infection. As shown in Figure 1, infectious chlamydial elementary bodies (EBs) were recovered from the UGT of the infected mice, thus demonstrating that *C. muridarum* reaches the tissue encompassing the oviduct and uterine horn as early as 3 days post-infection. We consistently recovered significantly more infectious chlamydial progeny from TLR3-deficient mice at all time points tested in each individual experiment. These findings are consistent with our previous reports of more robust *Chlamydia* replication in the absence of TLR3 function [6, 8, 27]. The observation that there was significantly more chlamydial progeny in the UGT tissues of mice early during the course of infection supports our hypothesis that TLR3 deficiency leads to a more progressive infection, increases chlamydial spread, and demonstrates a more rapid ascension of *Chlamydia* into the upper reproductive tract tissues.

**Figure 1.**
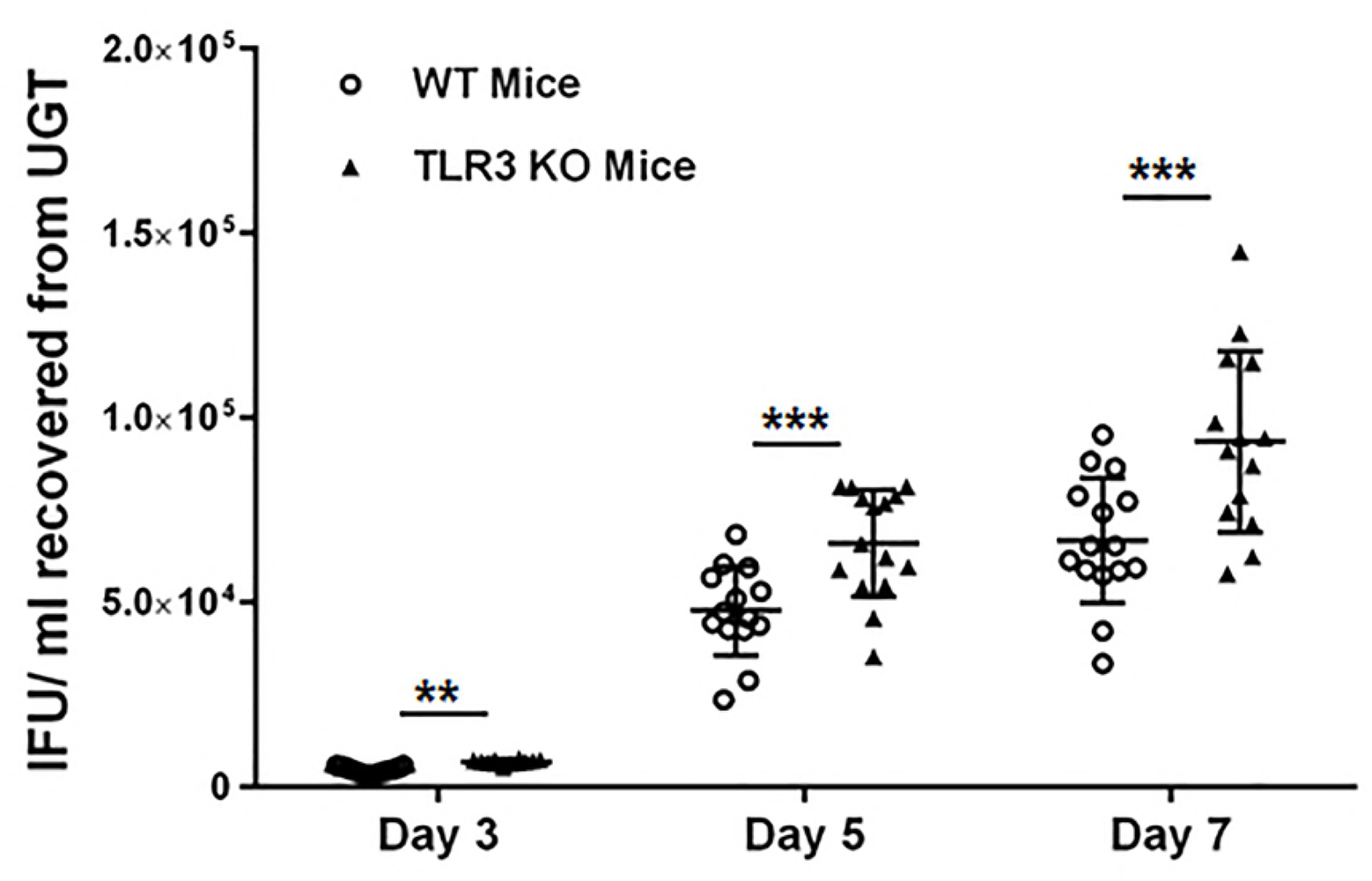
*C. muridarum* recovery from upper genital tract (UGT) tissue of wild-type and TLR3-KO mice. Genital tract infections were performed and UGT tissue was collected on days 3, 5, and 7 post-infection as described in Materials and Methods. The live chlamydial organisms recovered from the UGT were titrated, and the results were expressed as IFUs/ml recovered along the Y axis. Data shown are pooled results of 3 individual experiments resulting in the total of *n*=15 mice for each strain on days 3, 5, and 7. IFU= inclusion forming units. **= *p <0.005 and* ***= *p <0.001 when comparing WT vs. TLR3-deficient mice on that given day.*

### TLR3 deficiency augments the chlamydial degradation of OE cell TER

It has been demonstrated in other studies that *Chlamydia* infection can impact epithelial barrier function by inducing the secretion of various inflammatory mediators that are hypothesized to modulate the synthesis of host proteins that form cellular tight junctions (TJs) and adherens junctions (AJs) [34–37]. To determine if TLR3 deficiency has any impact on *Chlamydia’s* ability to affect host cell barrier integrity, we infected wild-type and TLR3-deficient OE cells with 1 IFU/ cell *C. muridarum* and measured the trans-epithelial resistance (TER) of the cell monolayers over a period of 48hrs in ECIS. As shown in Figure 2A, the TER in wild-type OE cells is significantly increased in response to *Chlamydia* infection after 6hrs post-infection when compared to infected TLR3-deficient OE cells and the mock controls. However, the TER begins to precipitously decline after about 24hrs (40% reduction in TER in the wild-type OE cells at the 48hr time point, *p< 0.05*). These data are in accordance with previous studies showing a reduction in TER at late time point during *Chlamydia* infection [34]. However, we did not observe an initial increase in TER in the TLR3-deficient OE cells as we had seen in the wild-type OE cells, which corroborates reports by others suggesting a critical role for TLR3 in increasing epithelial barrier function early during the epithelial repair process [38, 39]. The rate of degradation in TER was significantly higher in the TLR3-deficient OE cells, which resulted in an almost 80% decline in TER by 36hrs post infection. We previously reported that *C. muridarum*-induced synthesis of IFN-β in TLR3-deficient OE cells was significantly reduced, and that *Chlamydia* replication in the absence of TLR3-induced IFN-β was significantly increased in those cells when compared to wild-type OE cells [6]. Pretreatment of TLR3-deficient OE cells with 50U/ml exogenous IFN-β for 1hr prior to infection resulted in a significant reduction chlamydial progeny; however, pre-treatment with exogenous IFN-β had no significant impact on chlamydial replication in wild-type OE cells [8]. To examine whether IFN-β has impact on the more rapid decline of TER during TLR3 deficiency, we pretreated TLR3-deficient OE cells with 50U/ml of exogenous IFN-β 1hr prior to infection; however, the TLR3-deficient OE cells exhibited no significant change TER during infection when compared to the untreated TLR3-deficient OE cells (Figure 2B). The ECIS data show that TLR3 deficiency in oviduct epithelium led to a more rapid decline in TER during *Chlamydia* infection, and the rate of TER decline was not affected by the addition of exogenous IFN-β which is known to restrict *C. muridarum* replication in these cells.

**Figure 2.**
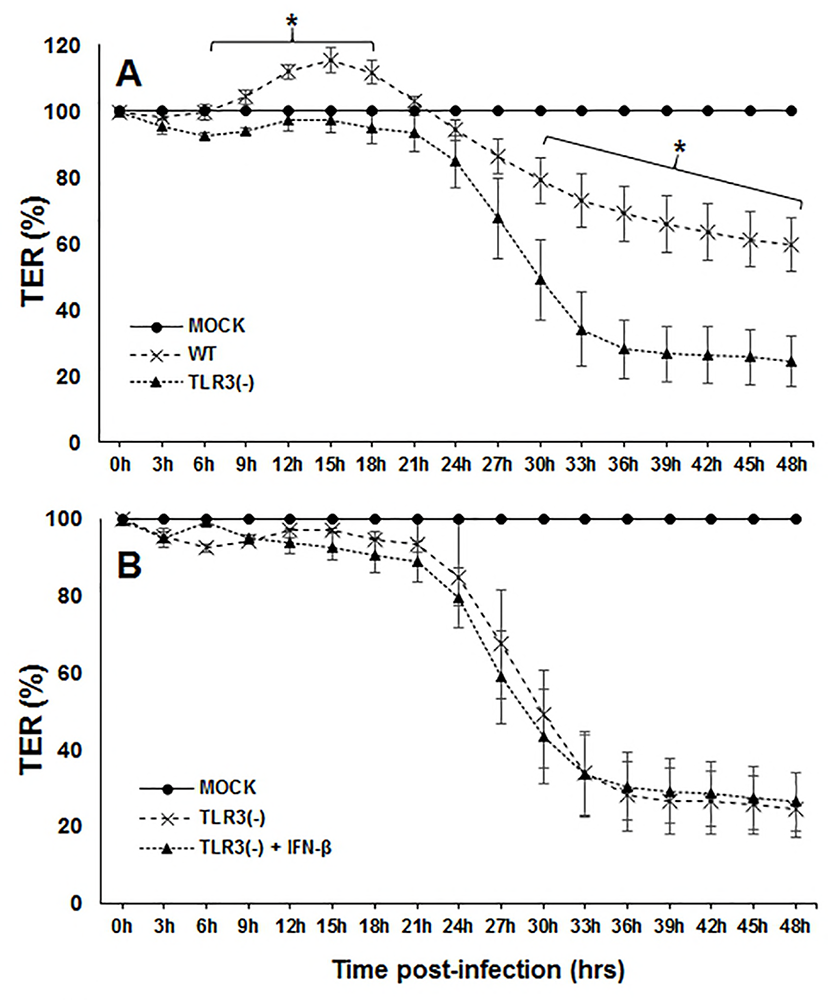
TLR3 deficiency exacerbates the *Chlamydia*-induced attenuation of trans-epithelial resistance (TER) in murine OE cell monolayers. TER was measured every three hours in *C. muridarum* infected: **A)** WT vs TLR3-deficient OE cells, and **B)** TLR3-deficient OE cells vs IFN-β pre-treated TLR3-deficient OE cells. TER at each time-point is relative to Mock-infected controls of each respective cell line set at 100%. Data are representative of 6 independent experiments. *= *p <0.05 comparing WT and TLR3-deficient OE cells at the given time.*

### TLR3 deficiency exacerbates the *Chlamydia* infection induced macromolecular permeability in OE cell monolayers

We next performed transwell macromolecular permeability assays to probe the physiological relevance of the pronounced decline of TER in TLR3-deficient OE cells during *C. muridarum* infection. Wild-type and TLR3-deficient OE cells were either mock-infected or infected with 1 IFU/ cell *C. muridarum* to determine if the drop in TER coincided with increased permeability of the OE cell monolayers to 70kDa FITC-dextran. As shown in Figure 3, *C. muridarum* infection significantly increased the amount of 70kDa FITC-dextran allowed to pass through the OE cell monolayers when compared to the mock-infected controls after about 18hrs post-infection in both cell types. However, the amount of FITC-dextran that traversed the TLR3-deficient OE cell monolayer was significantly higher than in wild-type cells after the 24hr time-point (*P< 0.05; denoted by asterisk*). We did not see any significant difference in the macromolecular permeability of the TLR3-deficient monolayer when the cells were pre-treated with 50U/ ml recombinant IFN-β for 1hr prior to infection (data not shown), indicating that IFN-β had little impact on the macromolecular permeability of TLR3-deficient OE cells during *C. muridarum* infection. The macromolecular permeability results corroborate the ECIS data demonstrating that TLR3 deficiency leads to more dramatic reductions in TER in murine OE cells. Collectively, the ECIS and transwell experiments indicate that TLR3-deficient OE cells are more susceptible to *Chlamydia* infection-induced breakdown of the monolayer integrity, and suggest that TLR3 signaling plays a substantial role in the maintenance of epithelial barrier function in oviduct epithelium during *Chlamydia* infection.

**Figure 3.**
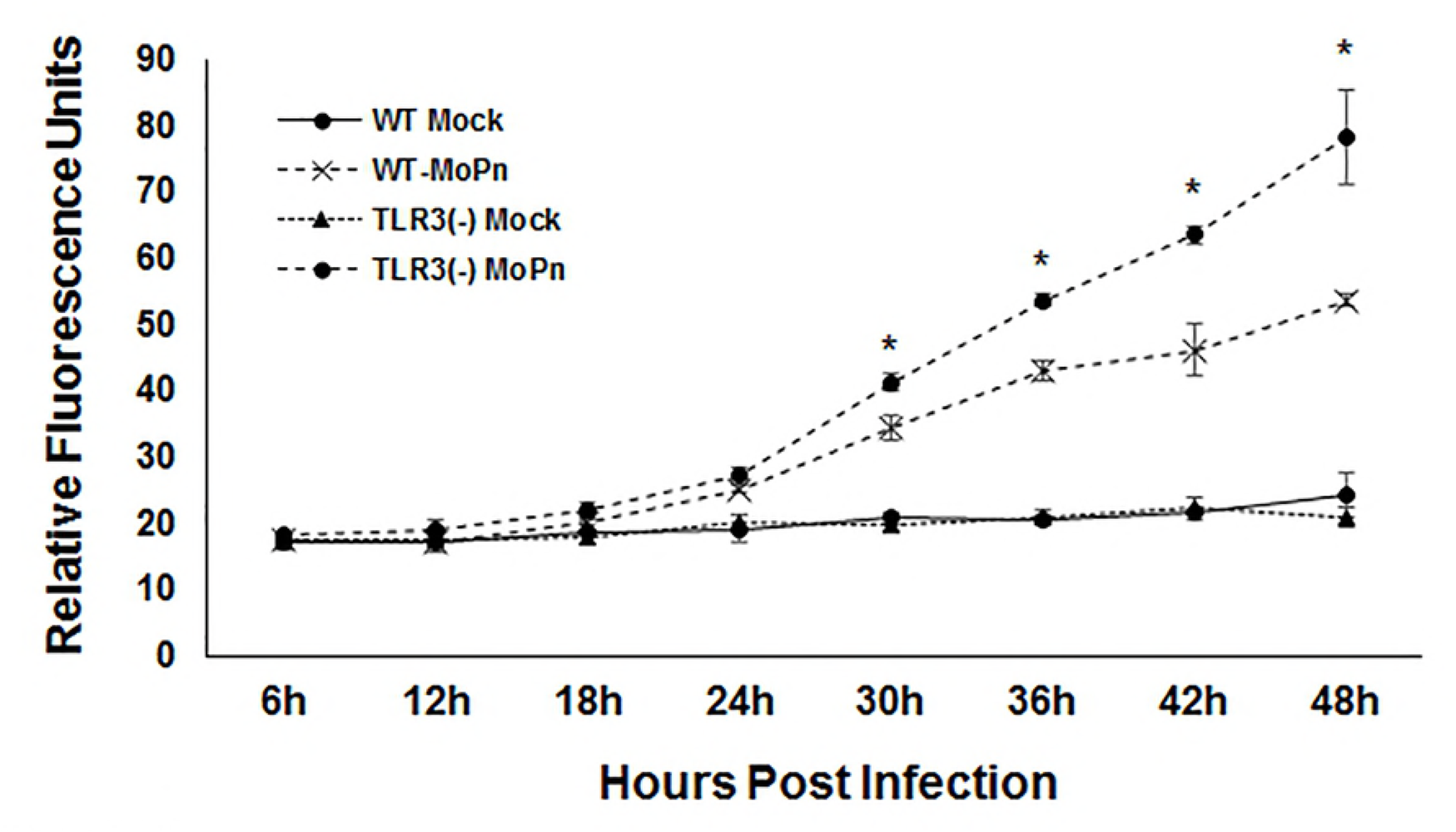
TLR3 deficiency leads increased rates of macromolecular permeability in OE cell monolayers during *Chlamydia* infection. Macromolecular permeability assays were performed in WT and TLR3-deficient OE cell lines that were either Mock-infected or *C. muridarum*-infected at a MOI of 1 IFU/ cell. Relative permeability was measured using a FITC-labeled dextran (70-kDa) probe. Samples were taken from the basolateral chamber of the transwell every 6hrs post-infection, and permeability was determined by increases in relative fluorescence compared to Mock-infected controls. Data are representative of three independent experiments. *= *p <0.05 comparing WT and TLR3-deficient OE cells at the given time.*

### The transcription of tight junction (TJ) genes are differentially regulated in TLR3-deficient OE cells during *Chlamydia* infection

The more dramatic reduction in TER appears to coincide with increased macromolecular permeability in the TLR3-deficient OE cells late during *C. muridarum* infection. This implies that TLR3-deficiency leads to a more acute breakdown in the integrity cellular tight junctions (TJs) and/ or adherence junctions (AJs) that are known to be affected during cellular invasion by certain viral and bacterial pathogens [34, 36, 37, 40–42]. To ascertain whether the dissimilarity in macromolecular permeability and TER between the wild-type and TLR3-deficient OE cells correlates with a differential regulation in TJ gene expression during *Chlamydia* infection, we measured mRNA expression levels of the candidate TJ proteins by quantitative real-time-PCR (qPCR). OE129-WT, OE-TLR3(-), and OE-TLR3(-) cells pre-treated with IFN-β were either mock-infected or infected with 10 IFU/ cell *C. muridarum* before being harvested for total RNA isolation at 0, 8, 12, 20, and 36hrs post-infection, and examined in subsequent qPCR for TJ gene expression. As shown in Figure 4, claudin-1 was significantly downregulated early (0-8h) and mid-infection (9-18h) in the OE129-WT cells, but eventually recovered to the levels of the mock-infected controls by 20hrs post-infection. However, claudin-1 gene transcription levels were significantly higher in the OE129-WT cells when compared to the mock-infected control cells by the 36hr time-point. In contrast, claudin-1 transcription was not as affected early during infection in the OE-TLR3(-) cells, and its expression levels were 3-fold lower than the OE129-WT cells at the 36hr time-point. Claudin-2 transcription was not very different between the 2 cell types, and its expression was also slightly downregulated early and mid-infection in both OE cell types. Claudins 3 and 4 gene expression levels were upregulated in both cell types; however, their expression levels were significantly reduced in the OE-TLR3(-) cells at most times past the 12hr time-point. There was no significant impact on transcription of any of the claudin genes tested when IFN-β was added to OE-TLR3(-) cells 1hr prior to infection, except for the 36hr timepoint when claudin-3 transcription was significantly higher in the IFN-β pre-treated TLR3-deficient OE cells.

**Figure 4.**
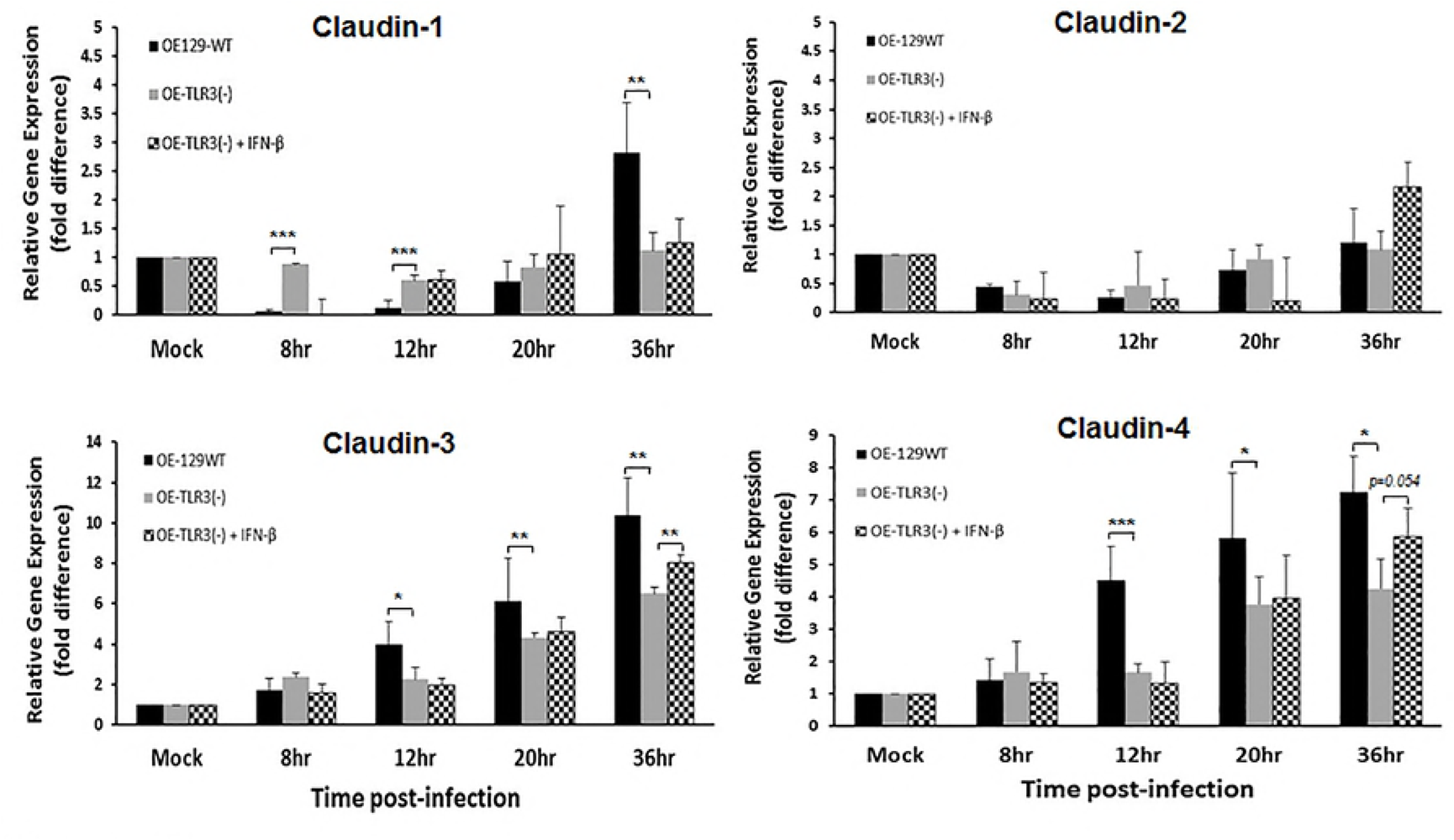
TLR3 deficiency dysregulates the *Chlamydia*-induced gene expression of the Claudin integral membrane tight junction (TJ) proteins. Gene expression levels of Claudins 1-4 were measured by qPCR at various times post-infection in *C. muridarum* infected WT, TLR3-deficient, and TLR3-deficient OE cells that were pre-treated with 50U/ ml recombinant IFN-β 1hr prior to infection. Data are representative of three or more independent experiments. *= *p <0.05*, **= *p <0.005, and* ***= *p <0.001 when comparing WT vs. TLR3-deficient OE cells at the given time.*

Figure 5 shows qPCR results of the *Chlamydia*-induced gene expression of ZO-1, JAM-1, and occludin in the OE129-WT, OE-TLR3(-), and IFN-β treated OE-TLR3(-) cells. As indicated, ZO-1 transcription levels were evenly downregulated during *C. muridarum* infection in all cell types, but gene expression levels began to rise in the OE129-WT cells starting at 20hrs post-infection. By the 36hr time-point, ZO-1 gene expression levels were more than 2-fold higher in the OE129-WT cells when compared to the OE-TLR3(-) cells. In contrast to ZO-1 results, we saw an initial uneven downregulation in JAM-1 transcription, but it was the OE-TLR3(-) cells that had a dramatic increase in JAM-1 gene expression starting at 20hrs post-infection. By the 36hr time-point, the JAM-1 gene expression was almost 8-fold higher in the OE-TLR3(-) cells when compared to the OE-129WT cells.

**Figure 5.**
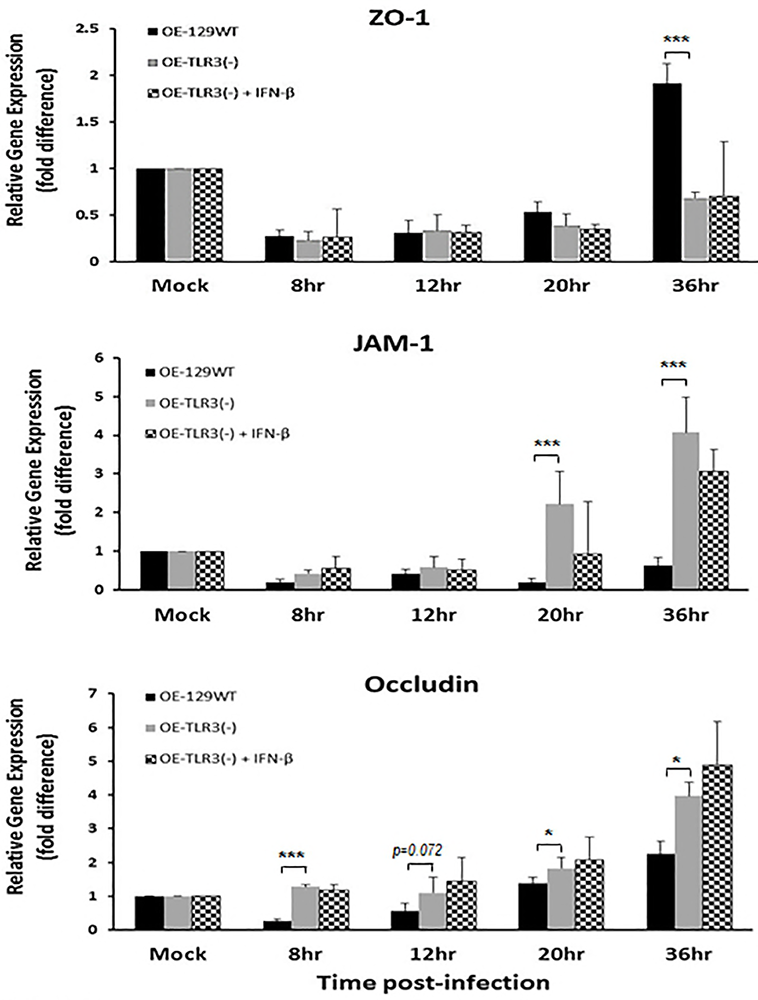
The *Chlamydia*-induced gene expression of the TJ proteins is impacted during TLR3 deficiency. Gene expression levels of ZO-1, JAM-1, and occludin were measured by qPCR at various times post-infection in *C. muridarum* infected WT, TLR3-deficient, and in TLR3-deficient OE cells that were pre-treated with 50U/ ml recombinant IFN-β 1hr prior to infection. Data are representative of at three or more independent experiments. *= *p <0.05 and* ***= *p <0.001 when comparing WT vs. TLR3-deficient OE cells at the given time.*

Occludin transcription was significantly downregulated early during infection of the OE-129WT cells, but slowly increased to a little over 2-fold difference in gene expression levels versus the mock infected cells at the 36hr time-point. In contrast, occludin transcription levels remained relatively stable in the OE-TLR3(-) cells before increasing to almost 4-fold versus the mock-infected controls at the 36hr time-point. Again, we saw no significant differences between OE-TLR3(-) cells and IFN-β treated OE-TLR3(-) cells. However, we did notice that the TJ gene expression levels trended towards the direction of their respective counterpart in OE-129WT cells in all cases except for occludin, where it trended more towards the OE-TLR3(-) cell results. Collectively, our data show that TJ gene expression levels in OE cells are affected by *Chlamydia* infection, which supports the findings of other investigators using different cell types [34, 35]. However, we show that the *Chlamydia’s* impact on TJ gene expression in OE cells are differentially regulated during TLR3 deficiency.

### TLR3 signaling has differential impact on the synthesis, stability, and cellular distribution of candidate TJ proteins during *Chlamydia* infection of OE cells

We next sought to determine if the *Chlamydia*-induced changes in the mRNA expression levels of candidate TJ genes translate into corresponding changes in their protein expression and/or stability. As shown in Figure 6A, ZO-1 protein expression levels in OE-129WT cells was rapidly diminished by 8hrs post-infection, was completely gone by 20hrs post-infection, but that there was little evidence of protein degradation. In contrast, ZO-1 expression was a bit more highly expressed in the OE-TLR3(-) cells, but appeared to be a lot less stable and showing more signs of protein degradation since there was no full-length ZO-1 protein at the 12hr time-point. Occludin protein expression was slightly upregulated early during infection in OE-129WT cells, but its expression level and protein stability appeared to remain steady throughout the course of infection. Occludin protein expression in the OE-TLR3(-) cells was slightly higher immediately after infection (0hr time-point) when compared the OE-129WT cells, but its expression was slightly down-regulated at the 8 and 12h timepoint before slowly increasing throughout the reminder of infection. The stability of the occludin protein in the OE-TLR3(-) cells was mostly unaffected by the *Chlamydia* infection as it was in the OE-129WT cells, and pre-treating the OE-TLR3(-) cells with IFN-β prior to infection had only minor impact on the stability and expression levels of both occludin and ZO-1.

**Figure 6.**
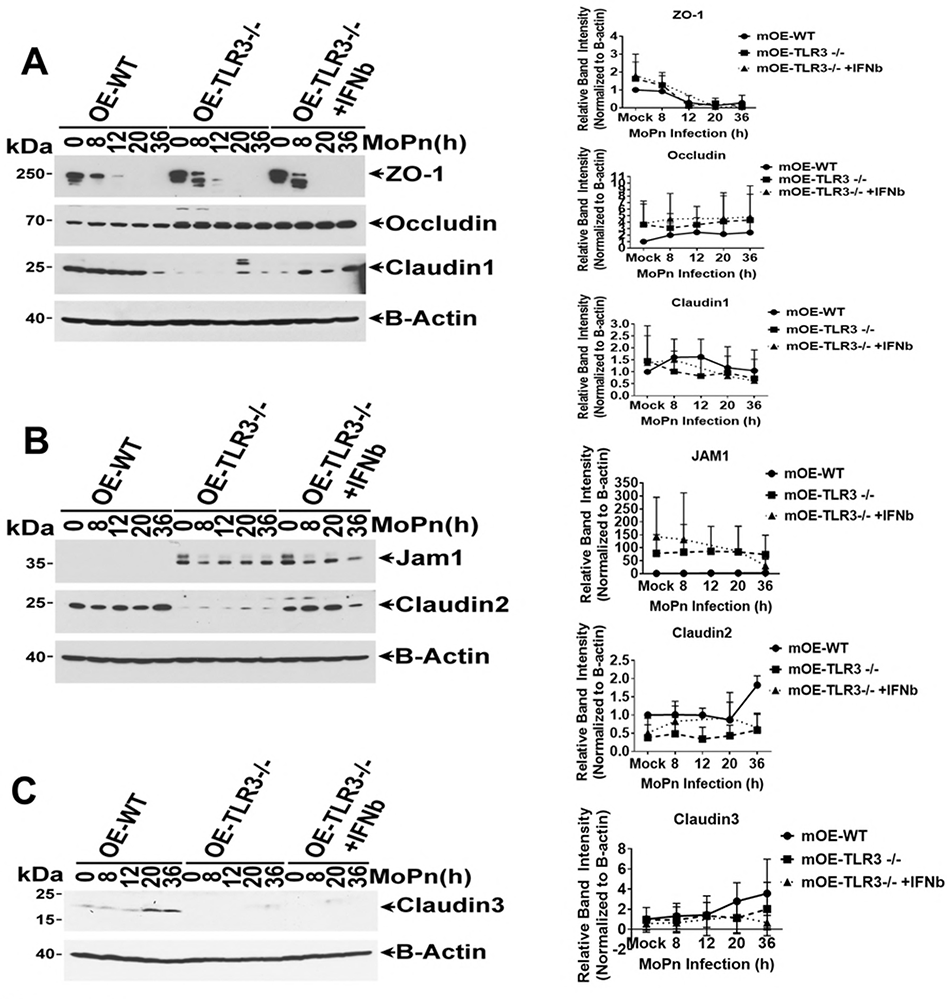
TLR3 deficiency leads to altered TJ protein expression in OE cell monolayers during *Chlamydia* infection. WT, TLR3-deficient, and IFN-β pre-treated TLR3-deficient OE cells were infected with 10 IFU/ cell *C. muridarum* for up to 36 hours before cell lysates were harvested at the various time-points shown. Protein expression levels were measured by western blot analyses for: **(A)** ZO-1, occludin, and claudin-1; **(B)** JAM1 and claudin-2; and **(C)** claudin-3. Right-side plots represent densitometry of tight-junction proteins normalized to β-actin control. Data are representative of three or more independent experiments.

The protein expression levels of claudins 1-3 were dramatically different between the OE-129WT cells and the OE-TLR3(-) cells (Figures 6A-6C). As shown, claudin-2 and claudin-3 proteins were expressed in the OE-129WT cells and their expression levels increased late during infection, while claudin-1 protein expression appeared to be diminished after the 20hr time-point. In contrast, the protein expression levels of claudins 1-3 were much lower in the OE-TLR3(-) cells; nonetheless, their expression levels were also moderately impacted by the *C. muridarum* infection. Interestingly, pre-treatment of OE-TLR3(-) cells with IFN-β prior to infection appeared mildly increase the synthesis and stability of ZO-1, while substantially bolstering the synthesis and stability of claudin-1 and claudin-2. These findings suggest that the *Chlamydia*-induced IFN-β synthesized via TLR3-signalling pathways plays a role in the regulating the protein synthesis of these genes, while also simultaneously enhancing their protein stability within the cell. In contrast to the claudin proteins, JAM-1 protein expression was substantially higher in the OE-TLR3(-) cells when compared to the OE-129WT cells where it was barely detectable on the immunoblot. Pretreating the OE-TLR3 cells with IFN-β prior to *C. muridarum* infection seemed to result in a diminished synthesis of JAM-1 over the course of *Chlamydia* infection, suggesting that the IFN-β synthesized via TLR3-signalling pathways plays a role in down-regulating the protein expression levels of JAM-1.

Fluorescent microscopy was also performed to assess protein expression levels and cellular distribution of various TJ proteins in WT and TLR3-deficient OE cells over the course of *Chlamydia* infection. As shown in Figure 7, we observed a differential distribution of JAM-1, ZO-1, and claudin-1 to the cellular TJs throughout the course of *Chlamydia* infection that seemed to depend on the functionality of TLR3. In corroboration with the western blot data, we saw substantially higher cellular expression levels of JAM-1 during *C. muridarum* infection in the TLR3-deficient OE cells, and its distribution along the outer membrane at cell-cell junctions increased at late times post-infection. In contrast, JAM-1 protein expression levels were noticeably lower in the OE-129WT cells and distribution at late times became increasingly cytoplasmic. ZO-1 protein expression levels along the outer membrane initially increased mid-infection in the OE-129WT cells, and its distribution remained mostly stable at cell-cell barriers late in infection despite the fact that its protein expression level had decreased. The expression and cellular distribution of ZO-1 differed during TLR3-deficiency in that its protein expression levels rapidly decreased over the course of infection, and its distribution became increasingly cytoplasmic in the TLR3-deficient OE cells late during *C. muridarum* infection. Claudin-1 protein expression levels remained mostly stable at mid-infection (12h) in the OE-129WT cells while only slightly increasing very late during infection; however, its distribution remained mostly stable at the cell-cell junctions throughout the course of infection. Claudin-1 protein expression levels seemed to slightly diminish at late times post-infection in the TLR3-deficient OE cells, and its cellular distribution remained mostly cytoplasmic throughout the course of infection. Results of the immunofluorescence studies reveal that *C. muridarum* infection can impact the cellular distribution of specific TJ proteins and likely the stability of cellular TJs of genital tract epithelium *in vitro*, which a finding that supports the investigation of others [34, 35, 43], and we demonstrate that TLR3 signaling plays some biological role in regulating this process. Collectively, our data implicates chlamydial infection, TLR3, and the IFN-β synthesized through TLR3 signaling pathways, in the assembly and stability of cellular tight junctions during *Chlamydia* infection of OE cell monolayers.

**Figure 7.**
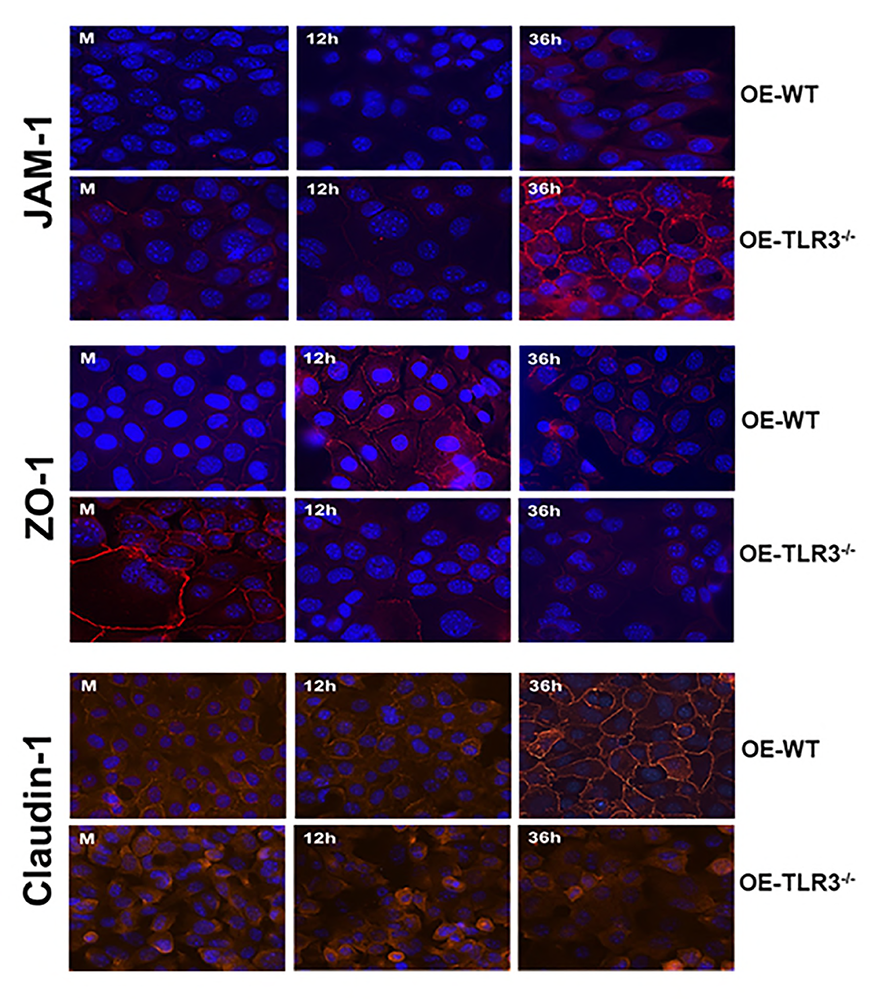
TLR3 deficiency impacts the chlamydial-induced rearrangement of Claudin-1, JAM-1, and ZO-1 in cellular tight junctions. WT and TLR3-deficient OE cells were infected with a low MOI of 1 IFU/ cell *C. muridarum* for up to 36 hours before cells were fixed at the listed time-point post infection and examined by immunofluorescence as described in **Materials and Methods**. The intensity of the TJ fluorescence in the infected cells was differentially regulated throughout the course of infection in the TLR3-deficient OE cells when compared to WT OE cells. All images are 40x magnification and nuclei were counterstained with DAPI (blue).

## Discussion

Our previous reports propose a protective role for TLR3 signaling in the immune response to *Chlamydia* infection in mice, and suggested that TLR3 signaling triggers mechanisms that bolsters the host’s ability limit the spread of *Chlamydia* and attenuate genital tract pathogenesis caused during infection. In this report, we investigated whether modulation of epithelial barrier function within the genital tract would be an example of such a mechanism where TLR3 signaling can exert its role in limiting *Chlamydia* spread and subsequent reproductive tract pathology. Our data showed that there were significantly higher amounts of *C. muridarum* in the UGTs of TLR3-deficient mice within the first 5 days of infection when compared to wild-type controls. These findings corroborate our earlier reports showing that TLR3 deficiency in mice leads to higher chlamydial loads in the LGT, but we now show that these mice also suffer from a more rapid ascension of *C. muridarum* into the UGT. The more rapid ascension of *Chlamydia* into the UGTs of TLR3-deficient mice is likely the combined result of dysregulation in the syntheses of multiple cytokines and chemokines that formulate the immunological barrier as we have previously shown [6, 8, 27], and a more rapid breakdown of physical barriers that functions to prevent the spread and transmission of sexually acquired infections [44, 45].

The major physical barrier in the female reproductive tract is the mucosal layer that lines the vagina, uterus, and luminal surfaces of the oviduct. The mucosal layer of the reproductive tract is an immunologically active layer that is comprised of an epithelial layer with tight cell-to-cell contact, which overlaid by viscous layer of mucus [45]. The impact of *Chlamydia* infection on the mucosal barrier function has been previously investigated by others, and it has been shown that *Chlamydia* infection disrupts the expression of critical proteins that form the cell-cell junctions (i.e., TJs and AJs) that play critical roles in forming intercellular adhesions [34–36, 46, 47]. The disruption of these intercellular adhesions during infection by *Chlamydia* is hypothesized to be an important immune-evasion strategy that involves recognition of specific chlamydial PAMPs, which then triggers signaling events that manipulate host-cell structure and function to help facilitate its spread within the host [43, 46]. In addition to the chlamydial downregulation in the secretion of specific immune regulators and anti-microbial peptides, the chlamydial disruption of intercellular adhesions is also thought to play a significant role in increasing host vulnerability to co-infection with other genital tract pathogens such as gonorrhea and HIV-1 [34].

The identification of specific cellular mechanisms that are involved in the disruption and subsequent destabilization of epithelial barrier function during *Chlamydia* infection is a recent area of study in the pathogenesis of genital tract *Chlamydia* infections. The chlamydial induction of the TLR-dependent inflammatory cytokines TNFα and IL-1β has been shown to up-regulate the expression of ZO-1 in murine endometrial epithelial cells, which results in significant changes in TER when compared to uninfected controls [35]. The *Chlamydia*-induced synthesis of TNFα and IL-1β via either TLR2 or TLR4 represents an example of such a mechanism in which the innate immune-response to *Chlamydia* infection leads to synthesis of inflammatory modulators that affect epithelial barrier function. IFN-γ is another TLR-dependent inflammatory mediator that is known to be induced during the protective immune response to *Chlamydia* infection [48–52]. Its synthesis in gut epithelium in response to either infection or inflammatory disorder has been shown to be associated with decreased TER and increased macromolecular permeability in gut mucosa [53]. Interestingly, we recently reported significant dysregulation in the syntheses of IFN-γ, TNFα, and IL-1β in *C. muridarum* infected TLR3-deficient mice when compared to wild-type controls [27]. Because IFN-γ, TNFα, and IL-1β are differentially expressed in the context of TLR3 deficiency, we hypothesized that TLR3 deficiency may have significant impact on the integrity of the epithelial barrier within the female reproductive tract during *Chlamydia* infection due to the altered expression of these and other key cytokines.

We showed that TLR3 deficiency does indeed have a significant impact on the TER of OE cell monolayers during *C. muridarum* infection, and the more rapid rate of TER decline in the TLR3-deficient OE cells resulted in the significantly increased rate in macromolecular permeability through these monolayers after 24hrs post-infection (see Figs 2 and 3). We also showed that TLR3 deficient cells were differentially regulated in the expression of several candidate TJ genes (Figs 4 and 5). The TJ protein expression in the OE cells appeared to be mostly representative of the gene transcription trends in the respective OE cell types. However, whereas we did not observe any significant impact on TER, macromolecular permeability, or TJ transcription levels when OE-TLR3(-) cells were pre-treated with IFN-β prior to infection, we did notice some augmented protein stability in ZO-1, claudin-1, and claudin-2 in the IFN-β pre-treated OE-TLR3(-) cells (Figs 6A and 6B). These findings indicate that the chlamydial induction of TLR3-dependent IFN-β during infection may play some role in the stabilization of the epithelial barrier, and proposes a possible mechanism of action that IFN-β uses to help limit the spread of *Chlamydia* as we have previously reported [8]. The chlamydial protease-like activity factor (CPAF) can degrade several host proteins including the adhesion protein nectin-1, which is a major component of cellular AJs [36, 37]. It is not yet known whether CPAF can target and degrade either claudin-1, claudin-2, or ZO-1, or even if CPAF activity can be regulated by IFN-β. However, based on our findings of protein instability of these particular TJ proteins, and the observed effect that IFN-β has on increasing their stability during *Chlamydia* infection, we can extrapolate that IFN-β may downregulate CPAF function as a possible mechanism of how TLR3-dependent IFN-β can help better control *C. muridarum* replication in wild-type OE cells and mice. Further study is needed to ascertain whether CPAF can degrade these particular TJ proteins over the course of *Chlamydia* infection, and to observe whether this activity is regulated by IFN-β.

Our data proposes a role for TLR3 signaling in maintaining the integrity of epithelial barrier function during genital tract *Chlamydia* infection, a function that we hypothesize is important in helping contain bacterial spread, but one that eventually becomes overwhelmed during productive infections as the pathogen ascends into the UGT. We show that TLR3 deficiency leads to more rapid ascension into the UGT of infected mice, and we attribute the more rapid ascension to dysregulation in the *C. muridarum*-induced syntheses of critical immune modulator and the infection caused disruption of cell-cell junctions. We are likely the first to report a role for TLR3 in maintenance of barrier integrity in oviduct epithelium during genital tract chlamydial infections; however, our work here corroborates a prior study in which the investigators reported a role for TLR3 in normal skin barrier repair following UVB damage [38]. In that study, the authors report changes to TER, paracellular transport of fluorescein-labeled sodium, and TJ protein expression when human keratinocytes were exposed to the TLR3 agonist poly-IC. The poly-IC was used to simulate the induction of snRNAs that are hypothesized to trigger TLR3 responses during UVB damage [39]. Our studies also parallel in that the prevailing hypothesis in both of our investigations involve a putative mechanism whereby TLR3 elicits this barrier maintenance function via cytokines and chemokines that are triggered upon stimulation by its appropriate PAMP. The identification of the exact immune factor that affect each specific aspect of the epithelial barrier function are subjects of our further investigation into this phenomenon.

## Acknowledgements

The authors thank Drs. Jerry Xu and David Nelson for their thoughtful critiques and their valuable scientific contributions towards the preparation of this manuscript. We also thank Stephanie Williams and Imani Crawford for their excellent technical assistance.

